# Immunoglobulins, MHC and T Cell receptors genes in Cetaceans

**DOI:** 10.1101/2020.10.24.353342

**Authors:** Francisco Gambón-Deza

## Abstract

Cetaceans correspond to mammals that have returned to the marine environment. Adaptive changes are very significant with the conversion of the limbs into flippers. It is studied the changes that have occurred in immunoglobulins, MHC class I and II and T cell receptors genes. Constant regions of immunoglobulins are similar to those of the rest of mammals. An exception is the IgD gene, which is composed of three CH domains but CH1 similar to CH1 of immunoglobulin M. In the IGHV locus, it exist a decrease in the number of VH genes with the absence of genes within Clan I. The number of V*λ* genes is greater than that of V*κ*. In the genes for T lymphocyte receptors, it exists a decrease in the number of V*α* genes with loss of significant clades and subclades. In V*β* and V*γ*, there is also the loss of clades. These declines of V*α*, V*β* and V*γ* are not present Artiodactyla, and they are specific to Cetaceans. In MHC present tree evolutive lines of class I genes. These species have DQ, DR, DO and DM genes, but they are no present DP genes.

## 1. Introduction

Cetaceans are mammals that returned to the aquatic environment. The closest terrestrial mammals are the Artiodactyla, more specifically the hippopotamus. In their adaptation process, gross phenotypic changes have occurred with the conversion of limbs to flippers and flukes. There are two parvordens. One is the Odontoceti (Cetaceans with teeth like the dolphin) and another called Mysticeti that encompasses the whales with a feeding system through filtering. Fossils of whale ancestors dating back 50 million years (Gatesy & O’Leary, 2001). In molecular studies, the divergence of the species present today began 35 million years ago (**?**).

The organs of the immune system have been studied in detail, finding no differences with terrestrial mammals. These studies strengthen the idea of the conservation of immune structures in mammals despite morphological and environmental changes in their evolution (Romano et al., 2002). However, this environmental change must impact the immune system. The presence of antibodies in serum was studied, and its messenger RNA sequenced (Nash & Mach, 1971; Andrésdóttir et al., 1987). An IgM sequence of RNAm is described in dolphin. This immunoglobulin in the transmembrane region shows a variation in the CART motif of uncertain significance (Lundqvist et al., 2002). In dolphins, two genes for IgG are described with characteristics similar to those of other mammals (Mancia et al., 2006). The sequence of an mRNA for IgA has also been published (Mancia et al., 2007) having similarity to the IgA of artiodactyls.

Of the genes of the T lymphocyte receptors, there is a study of the TRAV/DV and TRGV loci. They found a simple gene structure with few V genes in its loci (Linguiti et al., 2016). There is a recent study of the gene structure of MHC class II genes in cetaceans, concluding that they present a general pattern similar to that of other mammals (Zhang et al., 2019). There are also studies on the allelic variability of these proteins. Alleles are common in class I and DQB antigens. The presence of the same alleles in different species is interesting, suggesting a positive selection (Hayashi et al., 2003; Xu et al., 2009; Vassilakos et al., 2009).

In mammals, the number of V regions in the immunoglobulin and TCRs loci is variable (Olivieri et al., 2014b). There is currently no explanation for this variation. In that work, a low number of V genes evidenced in mammals that have evolved to adapt to an aquatic environment. Cetaceans are the most significant. In this work, it describes the characteristics of the genes of antibodies, V genes of TCRs and MHC of cetaceans.

## 2. Material and Methods

The entire study is with freely available data. The sequences of the genomes are in the NCBI, and on the web, https://vgp.github.io/. To obtain the sequences of interest, we use our applications developed in Python. For the general study of the sequences, the Biopython library was used (Cock et al., 2009). The exons of the constant regions of the heavy and light chains of the antibodies and the MHC class I and class II genes were performed with a machine learning (tensorflow (Abadi et al., 2016)) trained for their recognition. Other publications express the specific operation (Olivieri et al., 2020b). Vgeneextractor program similar to the previous one adapted for recognition of the V exons was used (Olivieri et al., 2020a). These applications extract the possible target exon. For this, a blast was performed with an artificial sequence that gives positivity to the objective. All hypothetical possible exons around the hit are translated into amino acids. The amino acid sequences are analyzed by machine learning, and the target sequences are obtained. Later, after a manual review, if the exons obtained are considered valid, they are added to machine learning.

The graphic representation of the exons was also done with a simple python application using the genomediagram library (Pritchard et al., 2006).

The amino acid sequences obtained were aligned with the MAFFT program (Katoh et al., 2005). The trees were built with the Fasttree program (Price et al., 2010) using the LG matrix (Le & Gascuel, 2008) and gamma parameter. The tree visualization was done with the Figtree program (http://tree.bio.ed.ac.uk/software/figtree/)

## 3. Results

Cetaceans are Laurasatheria mammals that have adapted to the marine environment. These animals have undergone significant phenotypic changes. We decided to study the essential molecules of the adaptive immune system.

At the time of writing this article, there are 12 reference genomes deposited with the NCBI. With computer tools (CHfinder) we look for the exons that code for CHs of immunoglobulins.

### 3.1. immunoglobulins

All species have at least one gene for each of the immunoglobulin classes present in mammals (table 1), Gene duplication found for IgG. All cetaceans have IgD with three exons for CHs. It stands out that the CH1 of IgD is identified to the CH1 of IgM while CH2 and CH3 are as IgD.

**Table 1:**
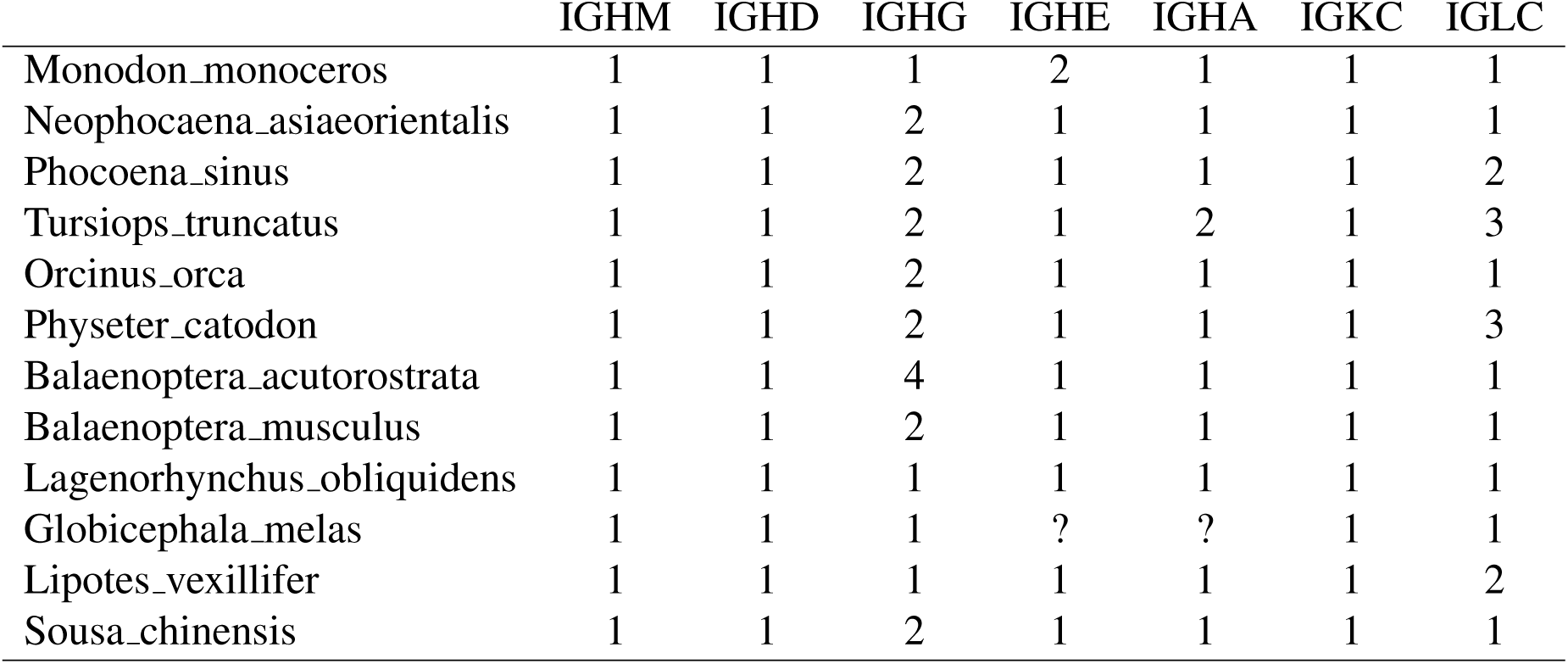
Number of genes for Immunoglobulins class

The study of 8 genomes is represented schematically in the figure 2. In general, all species have a very conserved position of genes with an IgM followed by an IgD, two genes for two isotypes of IgG, one gene for IgE and a gene for IgA. An exception is that of *Balaenoptera acutorostrata*, which has a gene for internal IgG in a standard position and three genes for 3 IgGs appear after gen for IgA. Also in *Physeter catodon*, additional exons of IgGs appear, but they must be part of pseudogenes.

**Figure 1:**
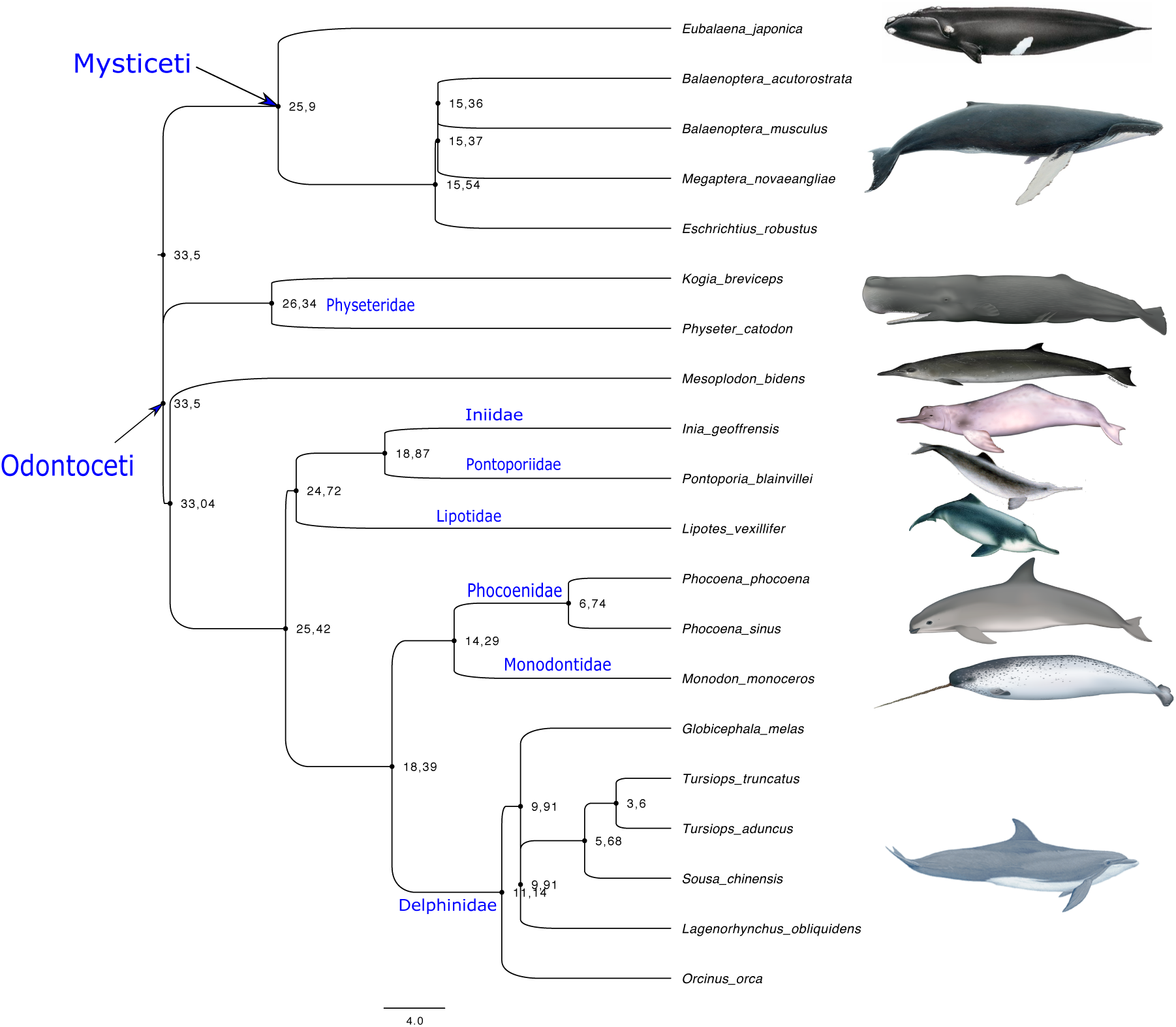
The figure represents the phylogenetic tree of the species present in this study. It has been obtained from the web http://www.timetree.org/. In the nodes, the divergence time is expressed.

**Figure 2:**
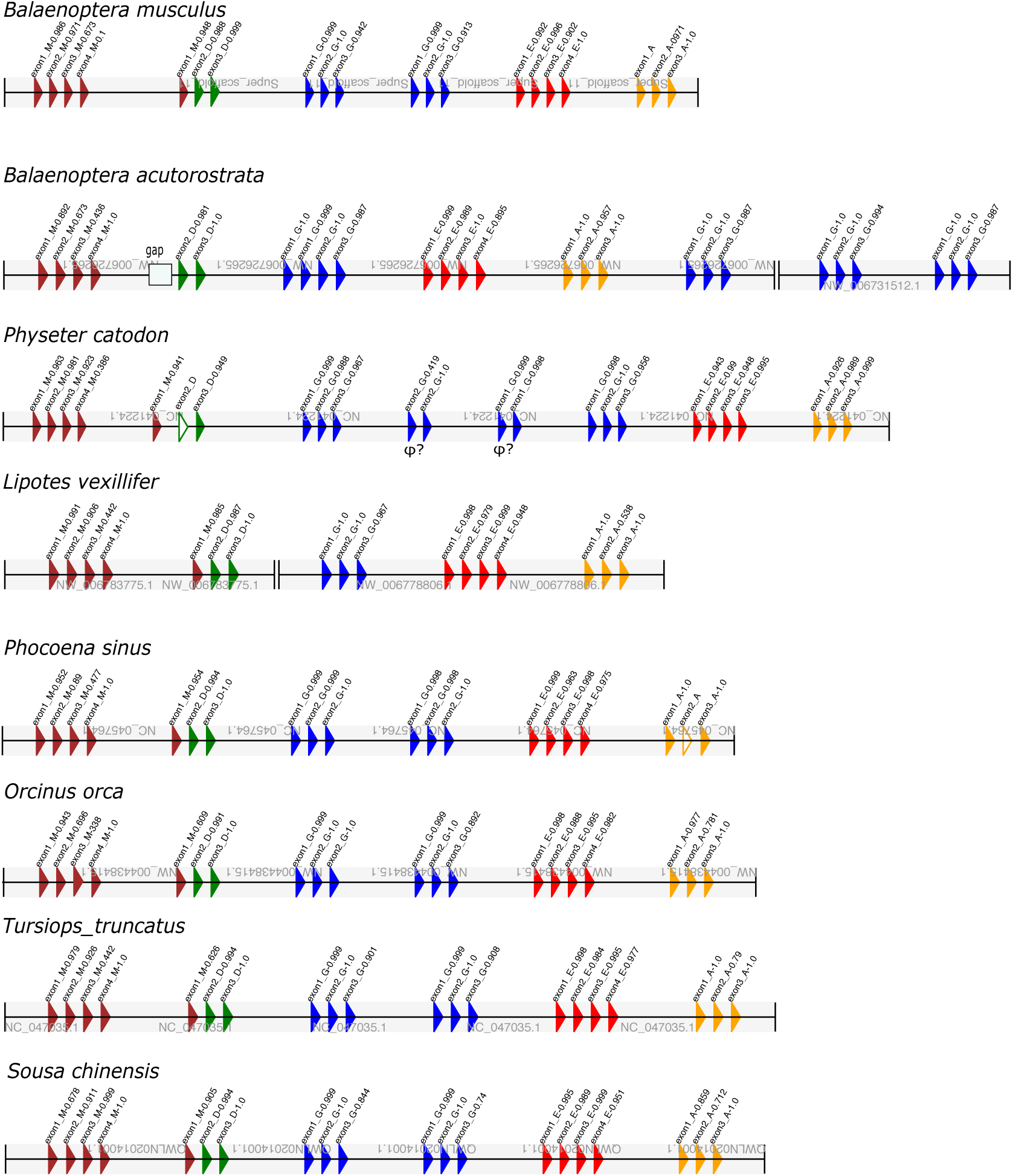
Schematic representation of the exons found and validated with the CHfinder application. In brown are exons identified as coding for IgM CHs. In green exons for IgD, in blue exons for IgG, in red exons for IgE and in yellow exons for IgA. In each exon, the numbering of the CH encoding is indicated. The probability of certainty is also expressed (1 is 100%). Exons not found with the CHfinder and that was found by the human review are unfilled.

The same CHfinder program has added training to identify the exons of the constant regions of the light chains of immunoglobulins. All species have one gene for the constant region of the kappa chain and 1 to 3 genes for the lambda chain (table 1).

### 3.2. V regions

Among the genes of immunoglobulins and those of T lymphocyte receptors, the genes that show the greatest changes in number enter species correspond to the V genes. We obtained the V exons of the seven loci that exist in mammals. The V exons obtained with the Vgeneextractor are expressed in Table 3 As can be seen, Cetaceans present a low number of V exons in all loci. Also highlights the presence of a greater number of V*λ* than V*κ* genes.

**Table 2:** Number of V genes in immunoglobulins and TCR loci

|  | IGHV | IGKV | IGLV | TRA/DV | TRBV | TRGC |
| --- | --- | --- | --- | --- | --- | --- |
| *Tursiops truncatus* | 47 | 4 | 37 | 15 | 8 | 2 |
| *Physeter catodon* | 8 | 6 | 16 | 18 | 12 | 2 |
| *Balaenoptera acutorostrata* | 6 | 7 | 18 | 17 | 12 | 3 |
| *Monodon monoceros* | 9 | 1 | 13 | 13 | 10 | 2 |
| *Lagenorhynchus obliquidens* | 5 | 3 | 17 | 9 | 7 | 2 |
| *Orcinus orca* | 26 | 2 | 19 | 7 | 8 | 2 |
| *Globicephala melas* | 11 | 7 | 13 | 10 | 7 | 2 |
| *Lipotes vexillifer* | 13 | 5 | 11 | 11 | 10 | 2 |
| *Neophocaena asiaeorientalis* | 15 | 6 | 23 | 9 | 11 | 2 |
| *Delphinapterus leucas* | 11 | 2 | 16 | 9 | 9 | 3 |
| *Phocoena sinus* | 10 | 5 | 10 | 11 | 11 | 2 |

**Table 3:** Number of MHC class I and II genes in cetaceans

|  | class_I | DPA | DPB | DQA | DQB | DRA | DRB | DMA | DMB | DOA | DOB |
| --- | --- | --- | --- | --- | --- | --- | --- | --- | --- | --- | --- |
| Tursiops.truncatus | 7 | - | - | 1 | 1 | 1 | 2 | 1 | 1 | 1 | 1 |
| Physeter.catodon | 3 | - | - | 1 | 1 | 1 | 1 | 0 | 1 | 1 | 1 |
| Balaenoptera.musculus | 3 | - | - | 1 | 1 | 1 | 2 | 1 | 1 | 1 | 1 |
| Monodon.monoceros | 5 | - | - | 1 | 1 | 1 | 2 | 1 | 1 | 1 | 1 |
| Lagenorhynchus.obliquidens | 4 | - | - | 1 | 1 | 1 | 2 | 1 | 1 | 1 | 1 |
| Orcinus.orca | 5 | - | - | 1 | 1 | 1 | 2 | 1 | 1 | 1 | 1 |
| Globicephala.melas | 5 | - | - | 1 | 1 | 1 | 1 | 1 | 1 | 1 | 1 |
| Lipotes.vexillifer | 6 | - | - | 1 | 1 | 1 | 1 | 1 | 1 | 1 | 1 |
| Neophocaena.asiaeorientalis | 4 | - | - | 1 | 1 | 1 | 2 | 1 | 1 | 1 | 1 |
| Delphinapterus.leucas | 3 | - | - | 1 | 1 | 1 | 2 | 1 | 1 | 1 | 1 |
| Phocoena.sinus | 4 | - | - | 1 | 1 | 1 | 2 | 1 | 1 | 1 | 1 |

In previous publications in which the Immunoglobulin sequences in the dolphin were described, they indicated the possibility of a low number of VH genes (Lundqvist et al., 2002). With all the VH sequences obtained from the exons in the germinal line of the cetacean species, we made a phylogenetic tree. Additionally we add the consensus sequences of the three clans that are represented in primates and the sequences of the VH present in the cow.

The distribution of the clades indicates the presence of particularities in the sequences (figure 3). There are sequences belonging to Clan III, and there are no sequences associated with Clan I. There is a high heterogeneity between sequences close to Clan II. The presence of an additional clade with a large number of sequences not associated with the three known Clans stands out. In this clade are the 21 VH exons of the cow. Within this new clade, the program extracted sequences that present an additional conserved segment of more of 30 amino acids with abundant prolines and cysteines amino acids. These sequences have already been described in the cow (Deiss et al., 2019). Here we find that the same process of conformation of long CDR3 must occur in cetaceans.

**Figure 3:**
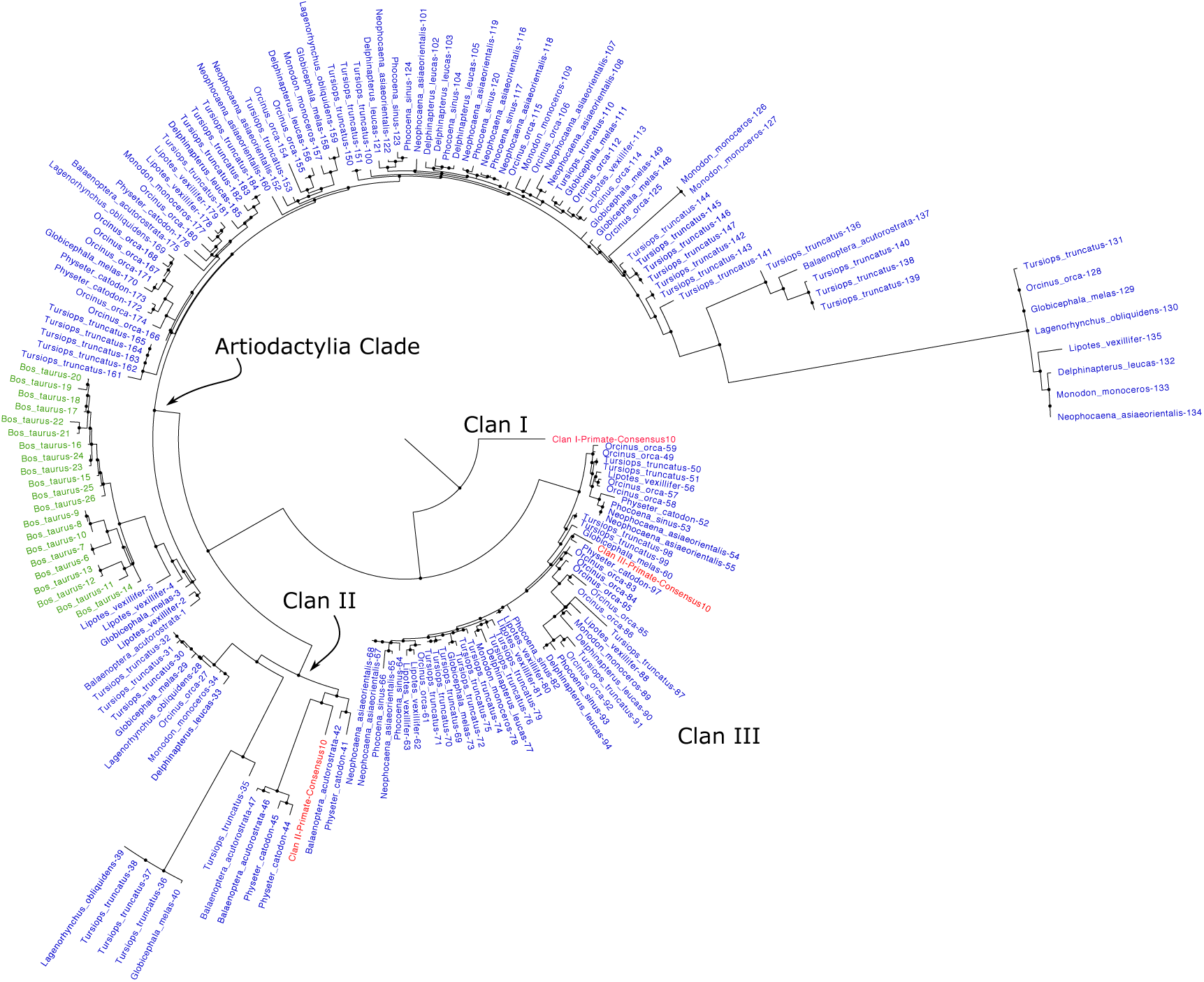
Figure present phylogenetic tree made with amino acid sequences obtained from the VH of cetaceans and cows. The consense sequence of each of the clans identified in the primates has been added. Taxa in red are sequences from primates, green from cow and blue from cetaceans. The alignment of the sequences was performed with the mafft program and the tree with fasttree using the LG matrix and gamma parameter. The visualization was made with the Figtree program.

Cetaceans for light chains have a higher number of V*λ* genes than V*κ*. Three V*κ* clans and at least 5 V*λ* clans have been described. the primate sequences were aligned with those obtained from cetaceans. As can be deduced from the phylogenetic tree presented in the additional figure 9 cetaceans have genes in each of the clans.

In previous publications, it was described in primates and rodents the presence of conserved V region sequences in the chains of T cell receptors (Olivieri et al., 2014c; Olivieri & Gambon-Deza, 2015). The sequences of V regions indicate probable evolutionary conservation of each one of them. That is to say, orthologous from each of the V regions exist in neighbouring species. Something different from what happens with the V regions of antibodies where the emergence of genes by duplication and subsequent selection occurs. These results suggest a probable determinism in recognition of the V regions of chains of TCRs, perhaps mediated by the need to recognize MHC molecules.

The deduced amino acid sequences of the V genes of the TRAV locus in primates and rodents conform six significant clades, which in turn harbour 35 subclades in total. The sequences found in the cetacean TRAV locus plus the consensus sequence of each primate clade were aligned. When constructing the phylogenetic tree with this alignment, the presence of the primate consensus sequences allows identifying if there are lines maintained with primates or if new ones are present.

The phylogenetic tree with the consensus sequences of each Primate clade in TRAV is in the figure 4. Four sequences outside the clades correspond to V*δ* sequences. All the others can be assigned to clades present in primates, indicating the conservation of these clades in evolution. Of the six main clades described in primates, II and IV do not have representatives in cetaceans. Of the 35 subclades found in the TRAV loci of primates, only 15 are present. The results demonstrate a selective decrease in the germinal repertoire of exons V. This diminished repertoire can be specific to Cetaceans or inherited from the common ancestor they share with Artiodactyla. To clarify this question, we obtained the V*α* regions of the cow genome. As can be seen in the additional figure 10 the cow has sequences rooted with most of the primate clades indicating its evolutionary conservation. Clades II and IV have representatives in the cow. These results indicate that V*α* genes losses occurred specifically in cetaceans.

**Figure 4:**
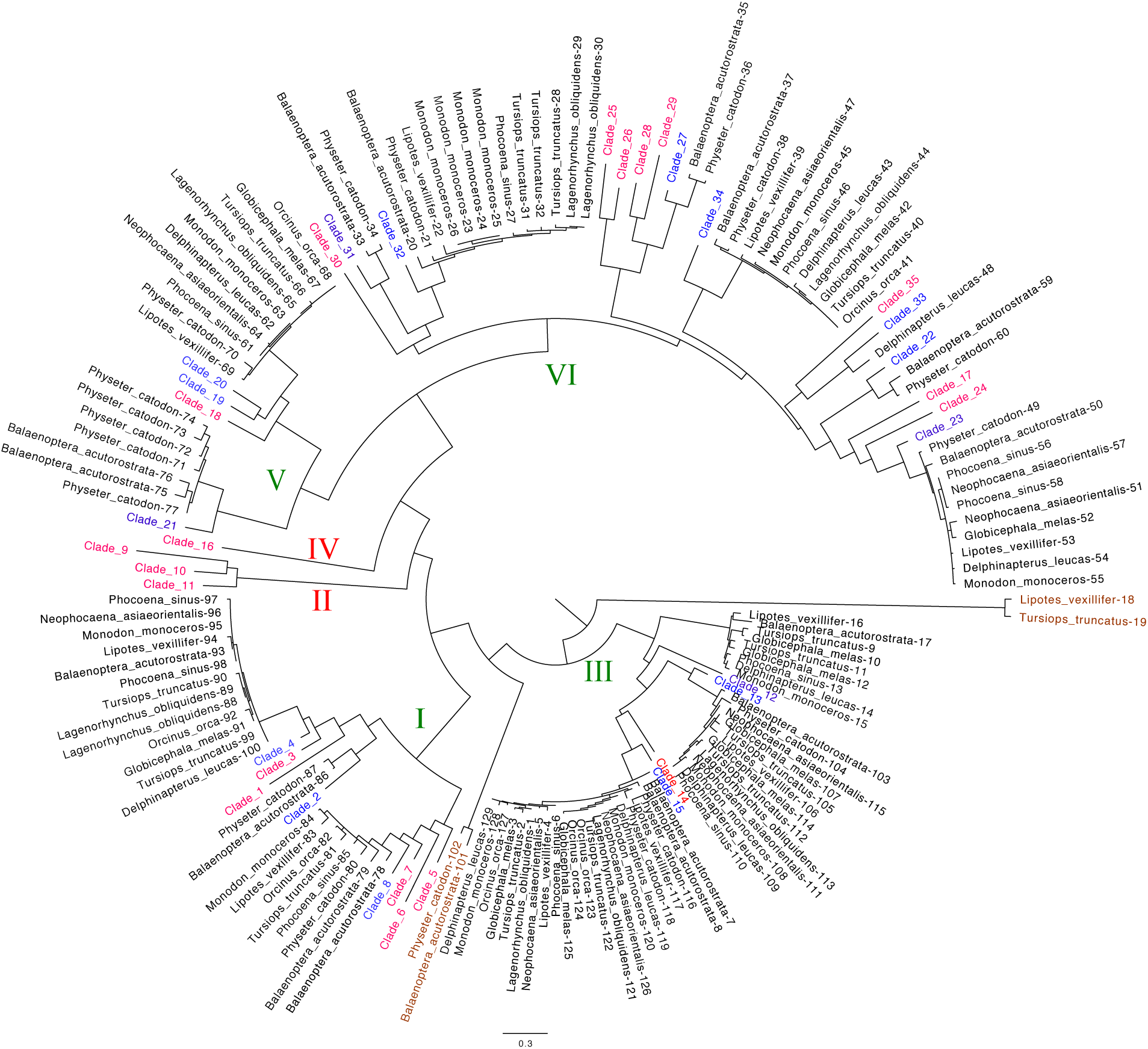
Figure present phylogenetic tree made with amino acid sequences obtained from the V*α* of cetaceans. The consense sequence of each of the subclades identified in the primates is represented. Taxa in red are consensus sequences from primates. Main clades in which they share primate and cetacean sequences are indicated in green roman numerals while those that do not have cetacean representation are in red. The alignment of the sequences was performed with the mafft program and the tree with fasttree using the LG matrix and gamma parameter. The visualization with the Figtree program.

The V*β* genes have an evolutionary pattern similar to that explained above for the V*α* genes. In primates and rodents, there are nine main clades. These main clades encompass a total of 25 subclades. The phylogenetic tree with the consensus sequences of each primate clade in TRBV is in the figure 5. There are some differences from the pattern described above for the TRAV. All the major primate clades except for clade III and IX have representatives in Cetaceans. The decrease in subclades of the TRBV loci concerning primates is smaller than in the TRAV loci (35 to 15 versus 25 to 15). In the case of cetaceans, two subclades within the main clade II are not in primates. As we did previously with the V*α* exons, we compared the V*β* exons of cetaceans with the V*β* from a cow (figure 11. Clades III and IX present exons in the cow, indicating that the loss is after the divergence of Artiodactyla. Also of the two new subclades found in cetaceans, only one of them is a related cow V*β*.

**Figure 5:**
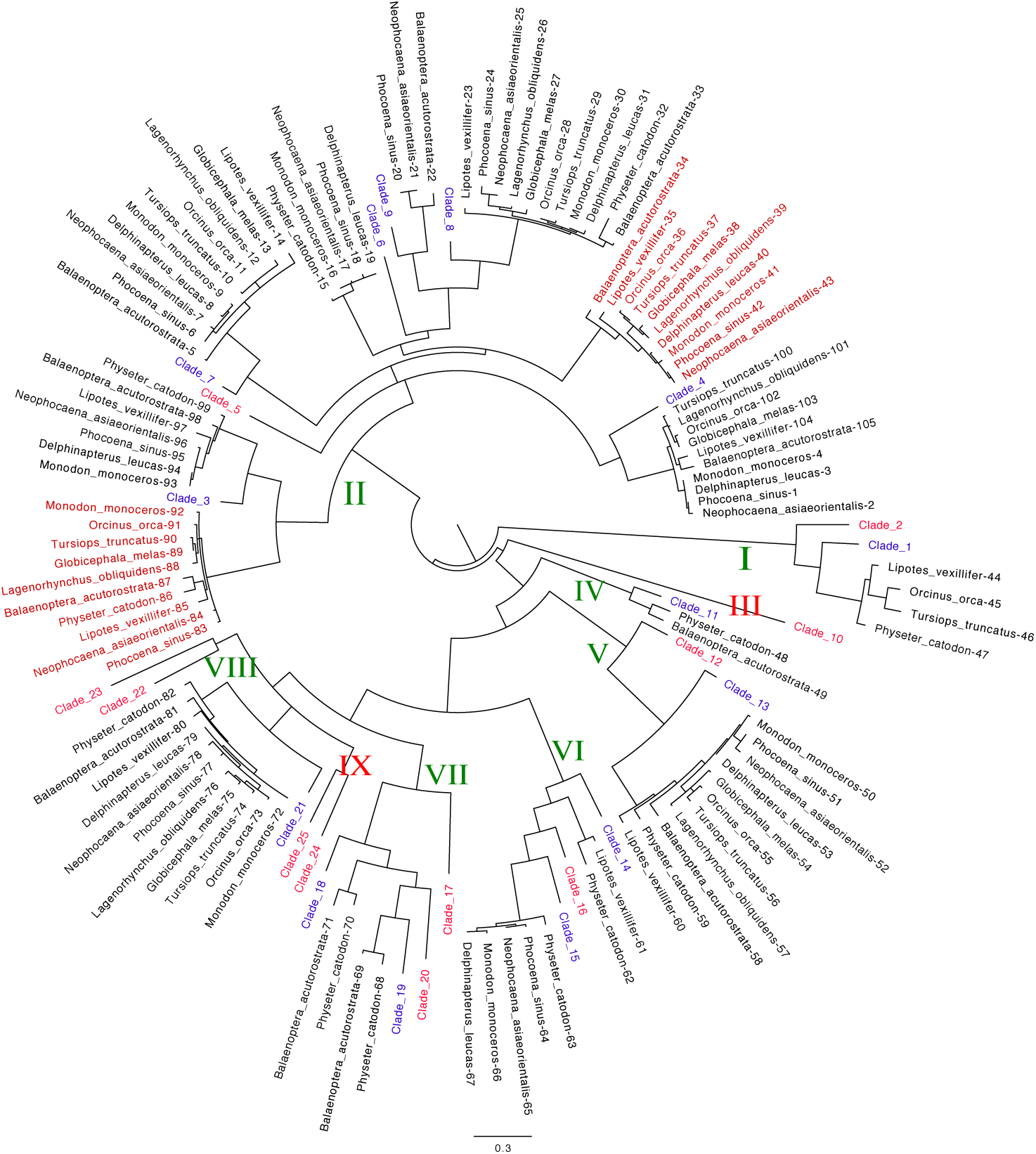
Figure present phylogenetic tree made with amino acid sequences obtained from the V*β* of cetaceans. The consense sequence of each of the subclades identified in the primates is represented. Taxa in red are consensus sequences from primates. Main clades in which they share primate and cetacean sequences are indicated in green roman numerals while those that do not have cetacean representation are in red. The alignment of the sequences was performed with the mafft program and the tree with fasttree using the LG matrix and gamma parameter. The visualization with the Figtree program.

With the Vgenextractor we also obtained the deduced amino acid sequences of the V*γ* exons. V*γ* genes in mammals are few in most species. In the phylogenetic trees carried out, they suggest sequence conservation. In figure 6, we have schematically represented the results found in cetaceans together with those found in cows and primates. The sequences obtained from the V*γ* exons are into three main clades. Clade II features two subclades. While primates and cows present sequences in all clades, cetaceans do not have sequences in Clade III.

**Figure 6:**
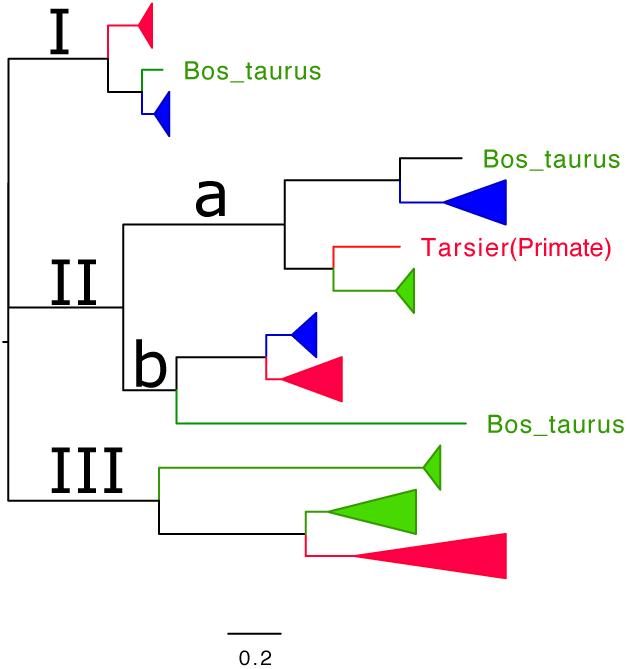
The figure represents the phylogenetic tree obtained from the alignment of V*γ* sequences of primates, cetaceans and cow. The clades are collapsed to reduce size. In red are the clades of primates, in blue of cetaceans and green of cow. The alignment of the sequences is with the mafft program and the tree with fasttree using the LG matrix and gamma parameter. The visualization is with the Figtree program

### 3.3. MHC class I and II

It searches for central exons of MHC class I and class II antigens in the genomes of cetaceans were with MHCfinder program. Clusters of exons 2, 3 and 4 of MHC class I and that of 2 and 3 of class II were probably viable genes. The appearance of loose exons is typical at these loci, and they are part of pseudogenes. In all species, the studied MHC genes are on the same chromosome. There is an exception of one isolated class I gene that is on another chromosome.

In class I antigens we detect three evolutionary (lines Line 1, 2 and 3, figure 8). Representatives of lines 1 and 3 are at one end with the class II genes. Line 1 has only one gene per species, while line 3 there are usually two or more per species. The genes of line 2 are on another chromosome, and there is one gene per species.

MHC class II antigens have already been studied by other authors (Zhang et al., 2019). The results with the CHfinder are consistent with those published. In this case, there are no exons with stop codons or with altered reading frames belonging to pseudogenes. Within the class two antigens, we evidenced the absence of DP sequences. The position of genes on the chromosome is in figure 7. From the end of the class I genes, a gene for the DR alpha chain appears first, followed most of the time by two DRB genes (these are in the chain complementary to that of the DRA gene). The DQA and DQB gene comes, the first being in the plus chain and the second in the minus, then the DOA, DMA, DMB and DOB genes appear successively. Of the genomes presented in figure 7, all present structure described above except for *Phisiter catodon* in which one of the DRB gene is lost, and the segment containing the DQ genes has undergone an inversion. Also, the DMA gene is not located.

**Figure 7:**
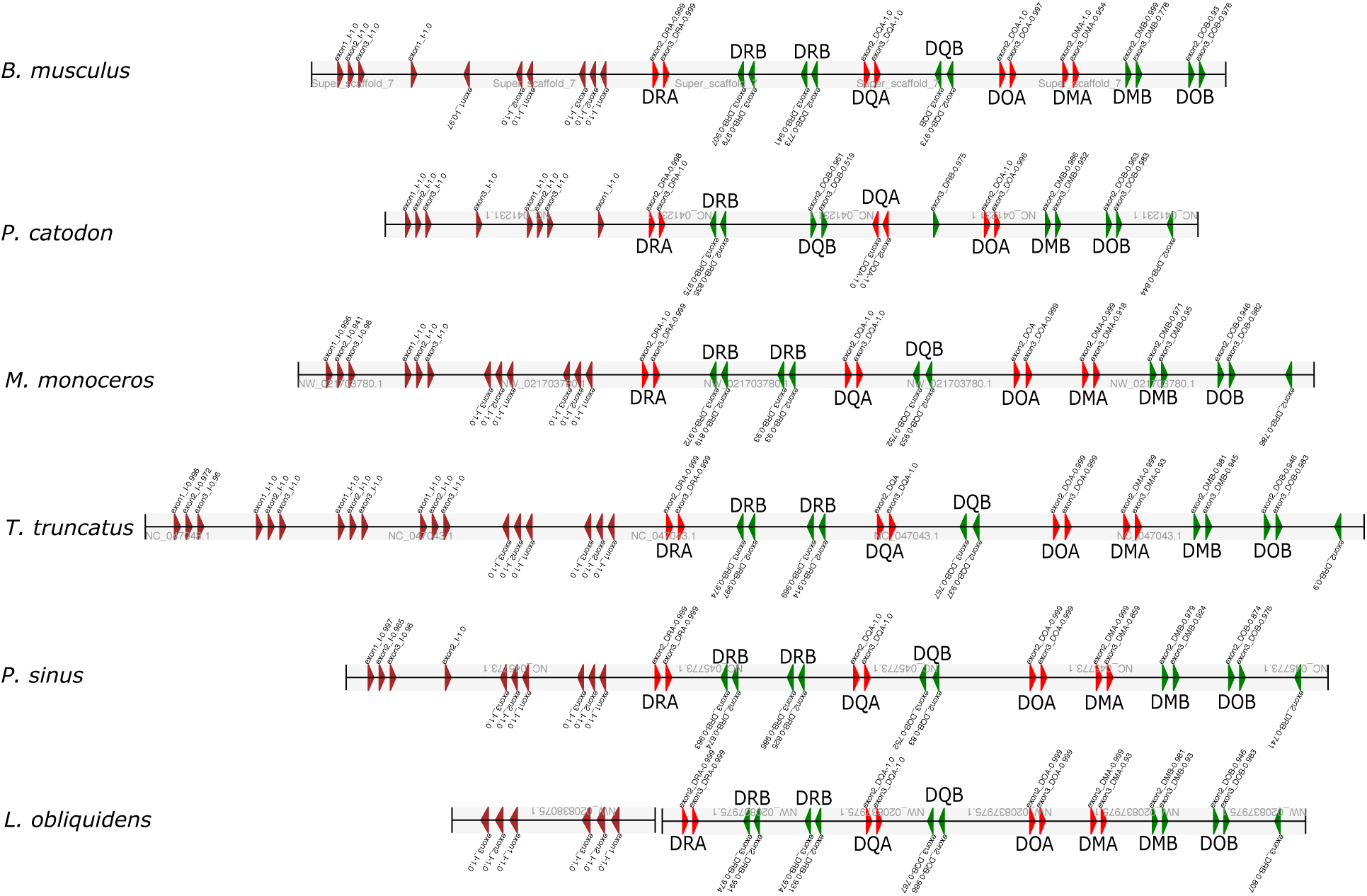
Schematic representation of the exons found and validated with the MHCfinder application. In brown are exons identified as coding for MHC class I antigens. In red exons for MHC class II alpha chain antigens and green exons for MHC class II alpha chain antigens. Each exon is numbered. The probability of certainty is also expressed (1 is 100%).

**Figure 8:**
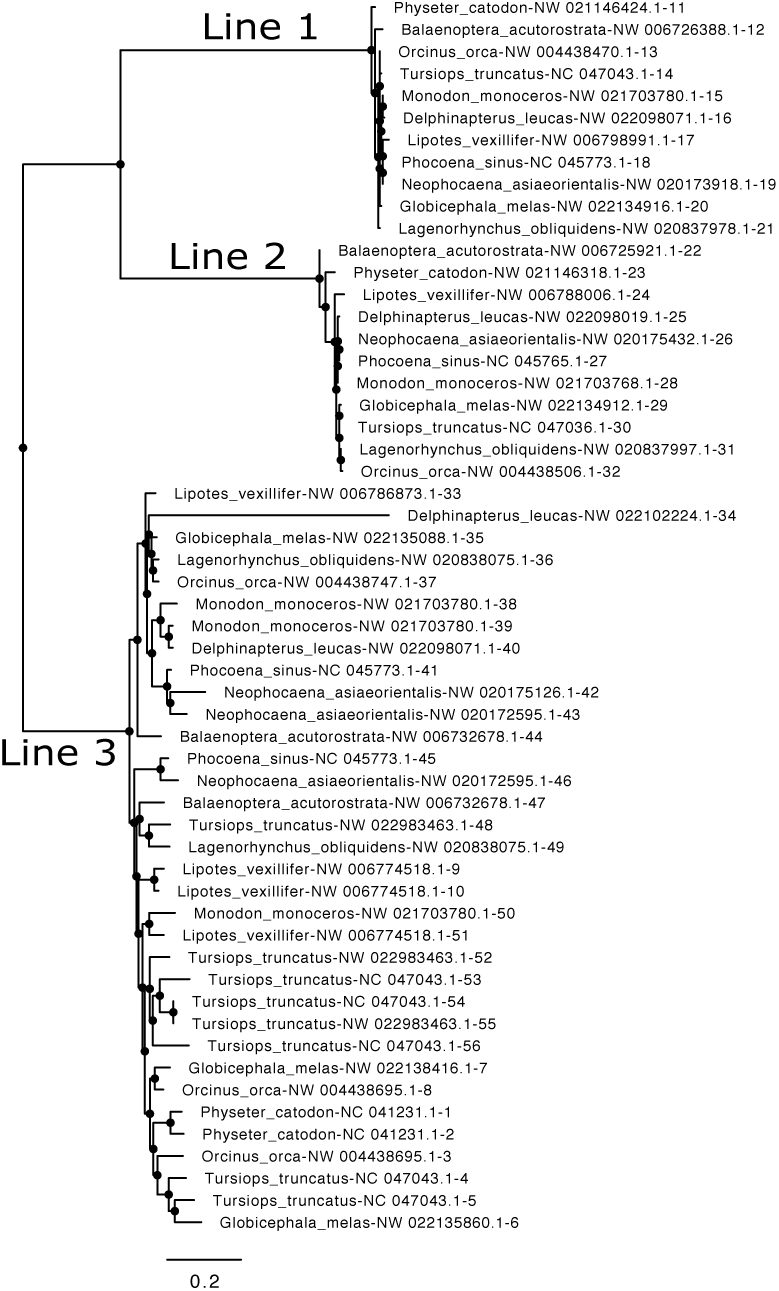
Phylogenetic tree of MHC class I found in cetaceans. The sequence of the antigens is from the sequences of the exons found with the MHCfinder application. The alignment is with the mafft program, the tree with the fasttree program (LG matrix, gamma parameter) and the visualization with Figtree.

**Figure 9:**
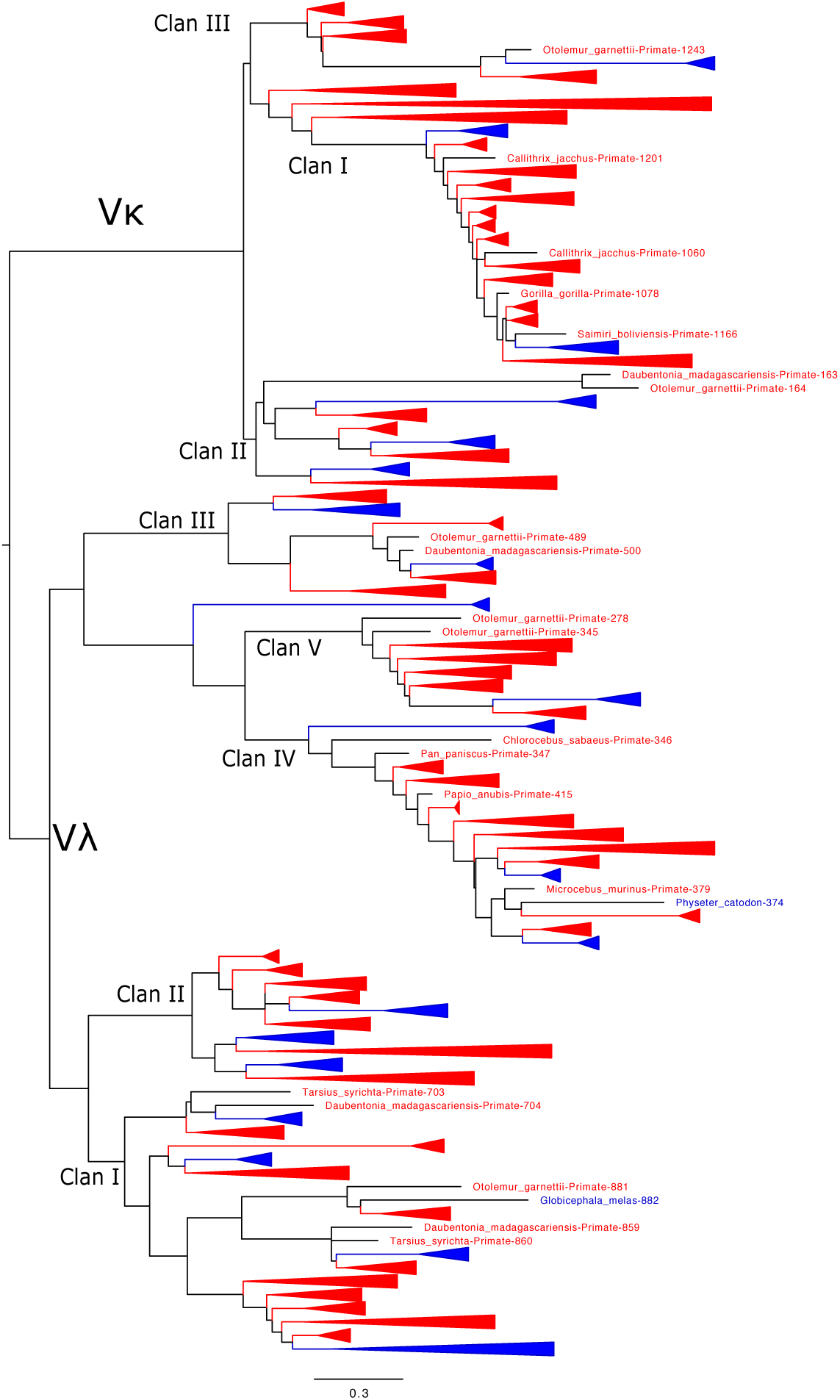
The figure represents the phylogenetic tree obtained from the alignment of V*κ* and V*λ* sequences of primates, cetaceans and cow. The clades are collapsed to reduce size. In red are the clades of primates, in blue cetaceans and in green cow. The alignment of the sequences is with the mafft program and the tree with fasttree using the LG matrix and gamma parameter. The visualization was made with the Figtree program.

**Figure 10:**
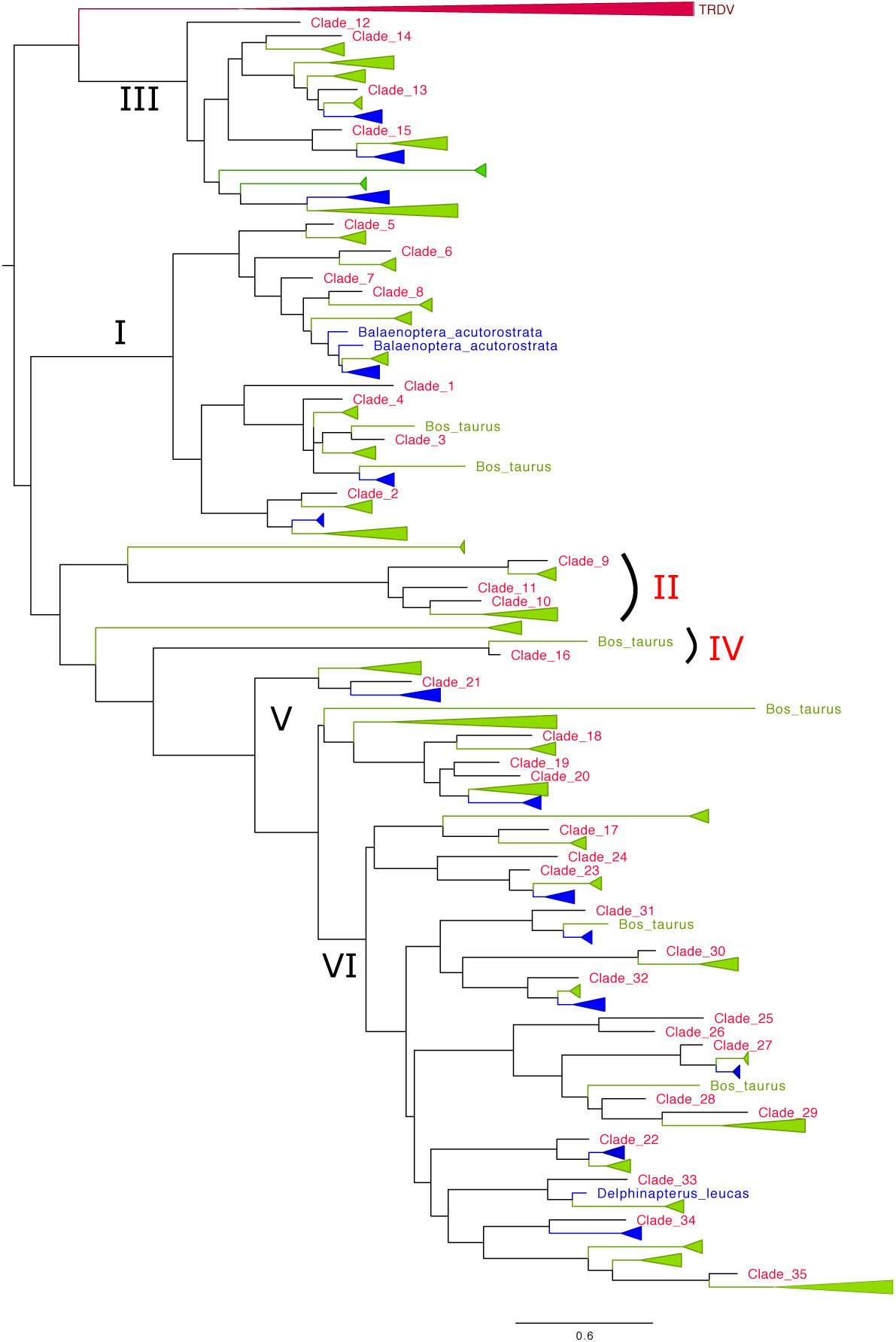
The figure represents the phylogenetic tree obtained from the alignment of V*α* sequences of primates, cetaceans and cows. The clades are collapsed to reduce size. In red are the clades of primates, in blue cetaceans and green cow. The alignment of the sequences is with the mafft program and the tree with fasttree using the LG matrix and gamma parameter. The visualization is with the Figtree program

**Figure 11:**
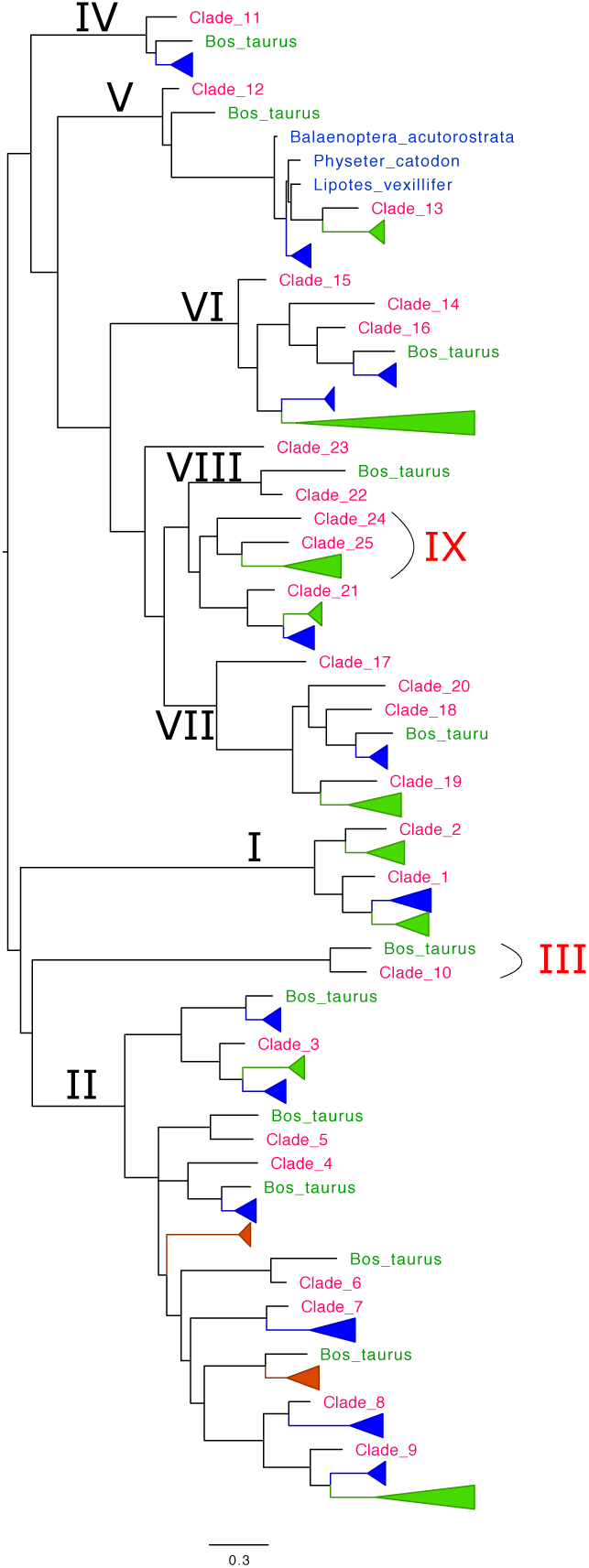
The figure represents the phylogenetic tree obtained from the alignment of V*β* sequences of primates, cetaceans and cows. The clades are collapsed to reduce size. In red are the clades of primates, in blue cetaceans and green cow. The alignment of the sequences is with the mafft program and the tree with fasttree using the LG matrix and gamma parameter. The visualization is with the Figtree program

## 4. Discussion

The study presented shows that cetaceans have constant regions genes from immunoglobulin and TCR genes similar to the rest of mammals. They also present histocompatibility antigens common to other mammals. The most significant differences are found in the number of V genes. An open test is why there are so many differences in the number of V genes in the germ line. Cetaceans are the mammalian species that have fewer genes than those described so far.

It remains unclear why two classes of light chains are necessary for mammals. The number of V regions is low. They have more genes for the lambda chain than for the kappa chain. Despite having a low number of genes, they have members in each of the clans described at the loci of these chains. Lambda light chains appear to have a different evolutionary process that kappa light chains. Various clans can be identified and appear to be conserved among species as separate as mammals and reptiles (Olivieri et al., 2014a). These results suggest a specific functionality of these clans that are not currently known. It is known that in terrestrial vertebrates, no species have existed that have lost the lambda chain, but several evolutionary lines have lost the kappa chain (snakes, birds) further supports this hypothesis. In this case, the decrease in V regions is more pronounced in kappa, again suggesting the biological importance of lambda chains.

The number of VH genes is low in all species. A specific loss is evidenced since no species has VH genes belonging to Clan I. This fact is inherited from an ancestor with artiodactylia as demonstrated by the absence in the cow. Some sequences present an elongation of the germ sequence of the exon for VH that conditions a very long CDR3 region with disulfide bridges. These VHs are in a clade that is difficult to assign to a specific clan and probably corresponds to a new clan associated with artiodactylia and cetaceans. There are sequences for Clan II and Clan III

Exciting results are those found in the number and distribution of exons for V*α* and V*β*. The presence of much fewer V*α* genes is evidenced, not presenting clades or subclades that are present in primates and rodents. Also, the V exons are related to specific primate clades. The absence of two main clades (Clades II and IV) at the TRAV loci is evident. Clade II in primates has three subclades indicating that it is a clade with a large presence of V genes. Clade IV of primates is unique since it is a clade that only has one V gene per species, suggesting a specific recognition. This V gene is absent in Cetaceans. Major clades are also missing at the TRBV locus. In this case, they are III and IX clades. It is noteworthy that clade III is a clade with one V*β* gene per species. Clade IV V*α* and clade III V*β* have similar evolutionary maintenance characteristics. The lost in cetaceans is suggestive that they may form a heterodimer with a specific function in primates. This function failed cetaceans, and therefore the clades have disappeared. If this were true, it leads to the assumption that there are selective pairings between V*α*s and V*β*s. There are human data on the presence of specific heterodimers shared between individuals (Tanno et al., 2020). With a greater number of studies in more species, it may be possible to confirm this hypothesis.

There is evidence that germline V*β* genes have predetermined affinity for MHC antigens. This evidence would support that the loss of V*β*s may be due to the loss of some MHC gene. Cetaceans do not provide conclusive data (Krovi et al., 2019). The MHC genes of cetaceans are similar to those of other mammals.

Is there a relationship between the V gene losses between the different loci? In previous publications, we have shown the existence of correlations between the number of V genes in loci between species (Olivieri et al., 2014c). The presence of a correlation between the number of V genes for immunoglobulins and TCRs is intriguing. In the adaptive immune response, the majority of antibody responses are T-dependent, suggesting the possibility of functional relationships between the V genes of the antibodies and the V*α* and V*β* genes of the TCR. New studies will be necessary to know if the loss of VH clade I in cetaceans is related to the loss of clades and subclades in the TRAV and TRBV loci.

## Supporting information

Sequeces exons

